# Immunization with SARS-CoV-2 nucleocapsid protein triggers a pulmonary immune response in rats

**DOI:** 10.1101/2021.08.24.457520

**Authors:** Everidiene K. V. B. Silva, Camila G. Bomfim, Ana P. Barbosa, Paloma Noda, Irene L. Noronha, Bianca H V. Fernandes, Rafael R. G. Machado, Edison L. Durigon, Sergio Catanozi, Letícia G. Rodrigues, Fabiana Pieroni, Sérgio G. Lima, Zelita A. J. Queiroz, Ives Charlie-Silva, Lizandre K. R. Silveira, Walcy R. Teodoro, Vera L. Capelozzi, Cristiane R. Guzzo, Camilla Fanelli

**Affiliations:** Renal Division, Department of Clinical Medicine, Faculty of Medicine, University of São Paulo, São Paulo, Brazil; Department of Microbiology, Institute of Biomedical Sciences, University of São Paulo, São Paulo, Brazil; Laboratório de Controle Genético e Sanitário, Diretoria Técnica de Apoio ao Ensino e Pesquisa, Faculdade de Medicina da Universidade de São Paulo; Laboratorio de Lipides (LIM-10), Hospital das Clinicas (HCFMUSP) da Faculdade de Medicina da Universidade de Sao Paulo, Sao Paulo, Brazil; Labinbraz Comercial Ltda. - Wiener lab, Brazil; Rheumatology Division of the Hospital das Clinicas da Faculdade de Medicina da Universidade de Sao Paulo - SP, Brazil; Department of Pharmacology, Institute of Biomedical Sciences, Universidade de Sao Paulo, Brazil

**Keywords:** SARS-CoV-2, COVID-19, vaccine, Nucleocapsid, immunization, antibody

## Abstract

The SARS-CoV-2 pandemic have been affecting millions of people worldwide, since the beginning of 2020. COVID-19 can cause a wide range of clinical symptoms, which varies from asymptomatic presentation to severe respiratory insufficiency, exacerbation of immune response, disseminated microthrombosis and multiple organ failure, which may lead to dead. Due to the rapid spread of SARS-CoV-2, the development of vaccines to minimize COVID-19 severity in the world population is imperious. One of the employed techniques to produce vaccines against emerging viruses is the synthesis of recombinant proteins, which can be used as immunizing agents. Based on the exposed, the aim of the present study was to verify the systemic and immunological effects of IM administration of recombinant Nucleocapsid protein (NP), derived from SARS-CoV-2 and produced by this research group, in 2 different strains of rats (*Rattus norvegicus*); Wistar and Lewis. For this purpose, experimental animals received 4 injections of NP, once a week, and were submitted to biochemical and histological analysis. Our results showed that NP inoculations were safe for the animals, which presented no clinical symptoms of worrying side effects, nor laboratorial alterations in the main biochemical and histological parameters, suggesting the absence of toxicity induced by NP. Moreover, NP injections successfully triggered the production of specific anti-SARS-CoV-2 IgG antibodies by both Wistar and Lewis rats, showing the sensitization to have been well sufficient for the immunization of these strains of rats. Additionally, we observed the local lung activation of the Bronchus-Associated Lymphoid Tissue (BALT) of rats in the NP groups, suggesting that NP elicits specific lung immune response. Although pre-clinical and clinical studies are still required, our data support the recombinant NP produced by this research group as a potential immunizing agent for massive vaccination, and may represent advantages upon other recombinant proteins, since it seems to induce specific pulmonary protection.

## INTRODUCTION

The disease caused by the new coronavirus Severe Acute Respiratory Syndrome Coronavirus 2 (SARS-CoV-2), known as COVID-19 (Coronavirus infection disease 2019) had it first reports in December 2019, in China, and by the end of January 2020, was defined by the World Health Organization (WHO) as a public health emergency of international importance. COVID-19 outbreak quickly reached virtually all the countries in the world, leading the WHO to change it classification, defining the disease as a pandemic [1]. Less than 2 years after the notification of the first COVID-19 patient, the world records more than 206 million of registered cases of the disease, 4,34 million of deaths caused by this pathology and the emergence of several SARS-CoV-2 variants [2–4].

The range of clinical presentations of COVID-19 is extremely wide and may vary from an asymptomatic condition, to severe respiratory and multiple organ failure, which may lead to death. The most common clinical symptoms include; dry cough, fatigue, loss of taste and smell, fever and dyspnea, which vary in intensity and may be moderate and comparable to the manifestations of a common cold, or severe, leading to the need for hospitalization and respiratory support due to acute lung injury [5]. In addition to the classic respiratory syndrome, other clinical and laboratory features, such as changes in the activation of blood coagulation pathways, were observed in COVID-19 patients, who may present fibrin deposition and disseminated microthrombosis, including in the nervous central system (NCS) and in the urinary system [6,7]. The pathophysiological mechanisms that determine the severity of COVID-19 manifestations are still unclear, and may depend on genetic susceptibility, pre-existing conditions; such as diabetes, hypertension and obesity, and individual general health. Main articles and case reports published to date, show the presence of at least two distinct phases in the evolution of COVID-19 pathology; The first, directly triggered by the viral infection, and the second, more severe and generally correlated with a worse prognosis for the patient, promoted by the exacerbated immune response of the host organism, a life-threatening systemic inflammatory syndrome known as “Cytokine Storm” [8,9].

The SARS-CoV-2 it the seventh member of the coronavirus family able of infecting human beings. In this family, the subtypes SARS-CoV (responsible for the severe respiratory syndrome outbreak in China, in 2003), MERSCoV (from the Middle East respiratory syndrome in 2012) and the new SARS-CoV-2 can cause serious illness in humans, while the subtypes HKU1, NL63, OC43 and 229E are associated with milder presentations [5,10]. SARS-CoV-2 is a ssRNA-virus, externally protected by a spherical-shaped phospholipid envelope of about 125 nm of diameter, covered by glycosylated Spike proteins (SP), which are the responsible for chemical affinity of the virus to the mammalian cells. SARS-CoV-2 binds the host cell through the interaction between its S protein and the transmembrane isoform of the angiotensin-converting enzyme 2 (ACE2) of host cells, that serves as a receptor, mediating viral entry. The SARS-COV-2 genome consists of approximately 30,000 nucleotides, which encode the structural viral proteins; S, Envelope protein (EP), Membrane protein (MP) and Nucleocapsid protein (NP), associated with protein units of NP, which in turn regulates viral replication replication [11]. SARS-CoV-2 seems to be less lethal than the SARS-CoV or the MERS-CoV, but more infectious, which could be contribute to its pandemic potential [12].

With the rapid spread of COVID-19 throughout the world, the development of vaccines against SARS-CoV-2 became necessary and urgent. Vaccines contribute to the development of immunological memory, thus minimizing the effects of infectious diseases, in a “second” exposition to the pathogen. Pathogen attenuation or inactivation, as well as the production of recombinant bacterial/viral-derived proteins are among the most employed biotechnology strategies for the production of vaccines [13]. Such elements stimulate both cellular and humoral adaptive immune response of the host, triggering the synthesis of specific antibodies against the pathogen, thus preparing it for future infections. Immunization through the vaccines is one of the most effective strategies for the prevention of infectious disease, and have being applied very successfully over the last decades, since it protects not only the patient who receives the immunizing agent, but the whole community, as the immunized person is unlike to become a vector of transmission to other people. The greater the number of people immunized, the lower the chances of a disease develops and becomes pandemic [13, 14].

Based on this assumption, research centers all over the planet, as well as pharmaceutical industries, public and private bodies have been working incessantly on the development of immunizers against SARS-CoV-2 since the beginning of 2020. Currently, a number of different vaccines against COVID-19 are already in emergency use, although the potential of immunizing action of each of these products, as well as the possible side effects that may be caused by them, have not yet been completely clarified. Moreover, in spite of the recent increase in the number of immunized people around the world, some countries are still suffering from the scarcity of available vaccines. Therefore, experimental and pre-clinical studies are still required to provide more details on the physiological mechanisms of immunization, and to contribute to the development of further immunizers against SARS-CoV-2 [13–15].

Taking the above into consideration, the aim of the present study was to verify the effects of the application of a recombinant protein derived from the viral NP of SARS-COV-2 virus, carried out by our research group through the culture of genetically modified bacteria, in 2 different strains of rats (*Rattus norvegicus*): Wistar and Lewis, thus verifying the safety of recombinant NP applications and the potential of this protein as an immunizing agent, by evaluating the production of specifics antibodies by sensitized animals.

## METHODS

### Cloning, Expression and Purification of SARS-CoV-2 Nucleocapsid Protein (NP)

SARS-CoV-2 RNA was isolated from the second Brazilian COVID-19 patient (GenBank: MT 350282.1) [16], and reverse transcription was performed to obtain the virus Nucleocapsid cDNA, which was used to amplified the nucleocapsid DNA fragment by Polymerase Chain Reaction, using the following primer sequences: 5’AGCATAGCTAGCTCTGATAATGGACCCCAAAATCAGC3’ (forward) and 3’ATTATCGGATCCTTAGGCCTGAGTTGAGTCAGC5’ (reverse). The amplicon DNA fragment was purified using GeneJET PCR Purification kit (ThermoFisher Scientific) and digested with BamHI and NheI FastDigest enzymes (ThermoFisher Scientific). The digested amplicon DNA fragment was cloned into the expression vector pET-28a 2, previously digested with the same pair of restriction enzymes, using T4 DNA Ligase enzyme (ThermoFisher Scientific). The DNA mixture was inserted into was used to transform E. coli STELLAR chemically competent cells (TaKaRa) and *E. coli Stellar* competent cells which were grown on 2xTY solid medium (16 g/L bacto-tryptone, 10g/l yeast extract, 5 g/l sodium chloride, 1.5 % agar) supplemented with kanamycin (50 μg/ml). Plasmid of positive clones were extracted using the GeneJET Plasmid Miniprep Kit following the manufacture protocol (ThermoFisher scientific). and tThe nucleocapsid cloning was confirmed by digestion with using BamHI and NheI FastDigest enzymes (ThermoFisher Scientific). The pET-28a containing the nucleocapsid DNA fragment was used for the expression of the recombinant protein in E. coli BL21 STAR (DE3) strain.

The transformed cells were cultivated in liquid 2xTY medium supplemented with kanamycin (50 μg/ml) and chloramphenicol (30 μg/ml). The transformed cells were grown until OD600nm of 0.6 was reached, under agitation of 200 rpm at 37 °C. At this stage, 0.5 mM isopropyl-β-D-1-thiogalactopyranoside (IPTG) was added for induction of nucleocapsid protein expression for 4 hours. The cells were harvested by centrifugation at 8,500 xg G for 15 minutes at 4 °C. The pellet was resuspended in lysis buffer (50 mM MOPS pH 7.5, 200 mM sodium chloride, 5% (v/v) glycerol, 0.03% Triton-100 and 0.03% (v/v) Tween-20) and lysed by sonication in a Vibracell VCX750 Ultrasonic Cell Disruptor (Sonics, Newton CT) in an ice bath under agitation. The lysate was centrifuged, at 30,000 xgG for 1 hour at 4 °C, and the supernatant was loaded to a HisTrap Chelating HP affinity column (GE Healthcare Life Sciences) previously equilibrated with 50 mM MOPS pH 7.0, 200 mM sodium chloride and 20 mM imidazole buffer. Bound proteins were eluted over 20 column volumes using 20–1,000 mM imidazole gradient. Samples of the eluted fractions were loaded into 15% SDS-PAGE gels and the fractions containing the nucleocapsid protein were concentrated using Amicon Ultra-15 concentric filters (Merck Millipore) with a 10 kDa cutoff. The concentrated sample was then loaded in a HiLoad 16/600 Superdex 75 pg column (GE Healthcare Life Sciences) for size exclusion chromatography previously equilibrated with 50 mM MOPS pH 7.0, 50 mM sodium chloride and 1mM Ethylenediaminetetraacetic acid (EDTA) buffer, and the protein eluted in a 50 mM MOPS pH 7.0, 50 mM sodium chloride and 1mM Ethylenediaminetetraacetic acid (EDTA) buffer. Samples of the eluted fractions were loaded into 15% SDS-PAGE gels and the fractions containing the nucleocapsid protein were The presence of the protein of interest in the solution was confirmed by Western Blotting (WB) and the fractions containing the protein were concentrated and stored at 4°C.

#### Animals and Experimental Protocol

All experimental procedures employed in the present study were approved by the Research Ethics Committee for the Use of Experimental Animals of the University of São Paulo Medical School (CEUA FMUSP N° 1522/2020). Adult male rats from Wistar and Lewis strains, weighing 250-300 g, were obtained from an established colony at the University of São Paulo. Animals were maintained at a constant temperature of 23±1°C, under a 12-12h light/dark cycle and had free access to tap water and conventional rodent chow.

Ten Wistar and 7 Lewis rats were subjected to intramuscular (IM) injections of 100μL of recombinant SARS-CoV-2 Nucleocapsid Protein (NP) diluted at 1,5 μg/μL in protein buffer, weekly, during 4 consecutive weeks: Groups Wistar NP and Lewis NP, respectively. Additional 5 Wistar and 5 Lewis animals received IM injections of 100μL of protein buffer (50 mM MOPS pH 7.0, 50 mM sodium chloride and 1mM EDTA) and were followed up during the same period, being used as control (Wistar Control / Lewis Control).

All rats had their body weight assessed twice a week and were kept in metabolic cages for 24h, weekly, to measure individual diet (g) and water (mL) consumption. Urine samples were also collected to determine 24h urinary flow (mL/24h) and urinary protein excretion concentration (UPE, mg/24h), using colorimetric methods (SENSIPROT #36 Labtest, Brasil). By the end of the study, after 4 weeks of follow up, rats were subjected to isoflurane inhalation anaesthesia and submitted to a xipho-pubic laparotomy. Blood samples were collected from abdominal aorta for biochemical measurements, while the left kidney, the liver, the brain and the lungs, were fixed in buffered paraformaldehyde, for further histological analysis.

#### Hepatic and Renal Function

Hepatic function was evaluated through biochemical dosage of proteins and enzymes in the serum samples of studied animals. Total serum protein was determined using Biureto LABTEST kit 99-1/250, serum albumin concentration was measured using Bromocresol green LABTEST kit 19-1/250, while the total Alanine Aminotransferase was measured in the serum rats using Liquiform-Cinética UV-IFCC LABTEST kit 108-4. Finally, Alkaline phosphatase were measured using Liquiform Bowers and McComb LABTEST kit 79-4.

Renal function analyses were performed in serum samples of the animals of each experimental group. Blood for serum obtaining was individually drawn from rats into tubes with clot activator and separating gel. Samples were submitted to 2,000 rpm centrifugation for 15 min, at room temperature. Blood Urea Nitrogen (BUN mg/dL) and serum creatinine concentration (Screat mg/dL) were assessed by colorimetric methods (UREIA CE #37 and CREATININA #35, respectively - Labtest, Brasil).

#### Plasma Lipid and Lipoprotein Analyses

Blood was individually drawn from rats into tubes containing anticoagulant and centrifuged for 20 min, at 3,000 rpm, at 4°C. The plasma was used for biochemical determinations as well as lipoprotein (LP) profile. Plasma total cholesterol (TC) and triglycerides (TG) concentrations were determined using enzymatic colorimetric kits (Labtest do Brasil). Plasma lipoproteins of Wistar and Lewis rats were fractionated and isolated using fast-performance liquid chromatography (FPLC) in a Superose 6 HR 10/30 column. One hundred microliters of plasma were applied to the column under a constant flow of 0.5 mL/min of Tris buffer (pH 7.0; 10 mmol of Tris, 150 mmol of NaCl, 1 mmol of ethylenediamine tetraacetic acid and 0.03% NaN3). Sixty fractions with 0.5 mL were collected automatically. TC and TG concentrations in the very low density LP (VLDL), low density LP (LDL) and high density LP (HDL) fractions were determined by enzymatic colorimetric kits (Labtest do Brasil) in a Cobas automatic analyzer (F Hoffman-La Roche, Basel, Switzerland).

#### Indirect Serum Biomarkers of COVID-19

C-reactive protein (CRP) and D-dimer were evaluated in serum samples using specific immunoturbidimetric test kits (Wiener Lab, Rosario, Argentina), while Troponin I (TnI) was determined in serum samples through a chemiluminescent immunoassay (Wiener Lab, Rosario, Argentina). All the analyses were performed in the SL 6000^®^ automatic analyzer (Wiener Lab Group).

#### Production of specific anti-SARS-CoV-2 antibodies by immunized animals

To verify the success of NP sensitization, we analyzed the presence of specific IgG anti-NP in the serum of inoculated animals by Western Immunoblotting (WB). All sera were previously treated with 5% skim milk with 10% of *E. coli* extract (supernatant of a lysed E. coli BL21 STAR (DE3) culture) for 2 hours, to block anti-*E. coli* antibodies from the solution, thus avoiding false-positive results. Meanwhile, purified recombinant NP samples were loaded to a 15% SDS-PAGE and then transferred to a nitrocellulose membrane of 0.2 μm cutoff using Tris-Methanol buffer (25mM TRIS-HCl, 192 mM glycine, 20% methanol, 0.1% SDS) in a Semi-Dry system (BioRad). The NP-containing membrane was then divided in test strips.

Each strip was initially blocked with 5% skim milk in Phosphate-Buffered Saline (PBS) buffer, 137 mM NaCl, 2.7 mM KCl, 12 mM Na2HPO4 pH 7.4, for 4 hours at 4°C, and then, incubated with the serum of each studied animal, diluted 1:1000 in PBS buffer, for 14 hours at 4°C. Membranes were then washed using PBS buffer containing 0.01% (v/v) Tween 20 solution, and then incubated with a secondary anti-Rat IgG peroxidase-conjugated antibody (Sigma-Aldrich), diluted 1:5000 in 5% skim milk, for 2 hours at 4°C. The reaction was developed with a Clarity Western Substrate kit (BioRad) and the image was captured in an Alliance Q9 chemiluminescence imaging system (Uvitec). Bands intensity was analyzed using ImageLab software (BioRad).

#### Histological Analysis

Samples from kidney, liver and lungs were dehydrated and paraffin-embedded through conventional techniques. “Blind” histological analyses of these organs were performed in 4-μm-thick sections, stained with hematoxylin-eosin.

For kidney tissue evaluation, 25 consecutive microscopic fields of renal cortex were observed under 200x magnification. We analyzed the presence of any glomerular and tubular architecture disruption, interstitial inflammatory infiltration and/or abnormal extracellular matrix accumulation.

Liver tissue analyses 25 consecutive microscopic fields of renal cortex were observed under 200x magnification. We analyzed the presence of any glomerular and tubular architecture disruption, interstitial inflammatory infiltration and/or abnormal extracellular matrix accumulation.

For lung histological evaluation, an immune cell infiltration score from 0 (no inflammatory infiltration) to 4 (100% of the microscopic field occupied by infiltrating inflammatory cells) was stablished. Lung analyses were carried up in 10 microscopic fields of peribronchiolar and interstitial area, under 400x magnification.

#### Immunohistochemistry of Lung Samples

Immunohistochemistry (IHC) for detection of macrophages (CD68+) and T-lymphocytes (CD3+) was performed in 4-μm-thick lung sections of each studied animal, in order to confirm and characterize the hypercellularity observed in the histological analyses. Specific immunostaining for macrophages was obtained by means of a streptavidin-biotin-alkaline-phosphatase IHC technique, using a primary monoclonal mouse anti-rat ED1 antibody (Serotec, #MCA341R, Oxford, UK), followed by a secondary biotinylated horse anti-mouse IgG antibody, rat adsorbed (Vector #BA2001, California, USA). CD68+ cells were developed in red, with a fast-red dye solution, and counterstained with Mayer’s haemalum (Merck, Darmstadt, Germany). For T-lymphocytes identification, immunoperoxidase staining was performed. The specific primary antibody polyclonal rabbit anti-human CD3 (Dako, #A4052, California, USA) was used, followed by polyclonal goat anti-rabbit IgG + HRP polymer (Spring Laboratories, #DHRR, California, USA). CD3+ cells were developed in brown, with a DAB dye solution.

Immunostained cells were quantified by histomorphometry. To access uniform and proportional lung samples, 10 fields (five non-overlapping fields in two different sections) were randomly analyzed in proximal and distal lung parenchyma. The measurements were done with a 100-point and 50 straight grid with a known area (103 μm^2^ at a 400x magnification) attached to the ocular of the microscope [17]. At 400x magnification, the number of positive immunostained cells in each field was calculated according to the number of points hitting positive cells for specific antibody as a proportion of the total grid area. Bronchi and blood vessels were carefully avoided during the measurements. In order to normalize the data, the number of positive cells, measured in each field, was divided by the length of each septum studied (to avoid any bias secondary to septal edema, alveolar collapse and denser tissue matrix seen in the fibrotic sections). The results were expressed as the number of positive cells/mm^2^ of connective tissue.

#### Statistical analyses

Results were presented as Mean ± SE. All analyses were performed using the GraphPad Prism 7^®^ software. Student’s t-test statistics was applied to compare the results obtained in groups NP vs. respective Control. Means were considered statistically different when p<0.05.

## RESULTS

### The immunization with recombinant NP was safe and well-tolerated by both Wistar and Lewis lineages of *Rattus norvegicus*

As exposed on **Table 1**, the 4 subsequent IM injections of 150 μg of recombinant NP were not toxic for Wistar nor Lewis strains of rats, since it was not observed any mortality among the experimental groups, and no one of the classic symptoms of physical suffering frequently observed in experimental animals, such as; reduced food or water intake, reduced mobility, impaired weight gain or breathing difficulties, were noticed in the animals included in the present study.

**Table 1.** General Parameters: Survival rate (%), Body weight (g), Dietary daily intake (g/24h) and Water consumption (mL/24h) of each experimental group.

| RAT STRAIN | GROUP | Survival rate (%) | Body weight (g) | Dietary intake (g/24h) | Water consumption (mL/24h) |
| --- | --- | --- | --- | --- | --- |
| WISTAR | Control | 100 | 367 ± 27 | 24 ± 3 | 40 ± 8 |
|  | NP | 100 | 374 ± 22 | 26 ± 3 | 33 ± 3 |
| LEWIS | Control | 100 | 331 ± 23 | 18 ± 1 | 16 ± 9 |
|  | NP | 100 | 339 ± 16 | 18 ± 2 | 16 ± 7 |

Hepatic and renal function of NP animals were carefully analyzed in order to investigate subclinical toxicity signs. Thus, even though the differences observed among the two strains of rats, the hepatic function panels of both Control and NP animals were compatible with the laboratorial reference ranges for healthy rats [18, 19]. As shown in **Figure 1**, in spite of statistical differences between Control and NP groups, all tested animals exhibited regular values of serum protein (**A**), albumin (**B**) ALT (**C**) and ALK (**D**) concentrations. Moreover, in order to further evaluate the hepatic metabolism of lipids in the animals submitted to NP injections, we analyzed plasma TC (**E**), cholesterol fractions (**F**), TG (**G**), and triglyceride fractions (**H**), which were all very similar among Control and NP animals of each studied strain. Finally, histological analysis of HE-stained liver sections, illustrated in **Figure 1**(**I**), showed no structural abnormalities in the hepatic architecture of both Wistar and Lewis NP animals, compared to their respective controls. There was also not observed any impact of IM injections of NP on the renal function of both Wistar and Lewis strains of rats. As can be seen in **Figure 2**, NP animals exhibited regular urinary flow (**A**), according to their lineage, did not presented proteinuria (**B**), nor Screat (**C**) or BUN (**D**) retention. Accordingly, no histological changes were observed in the renal parenchyma of animals submitted to NP injections, compared to their strain-matched controls, as illustrated in **Figure 2** (**E**).

**Figure 1.**
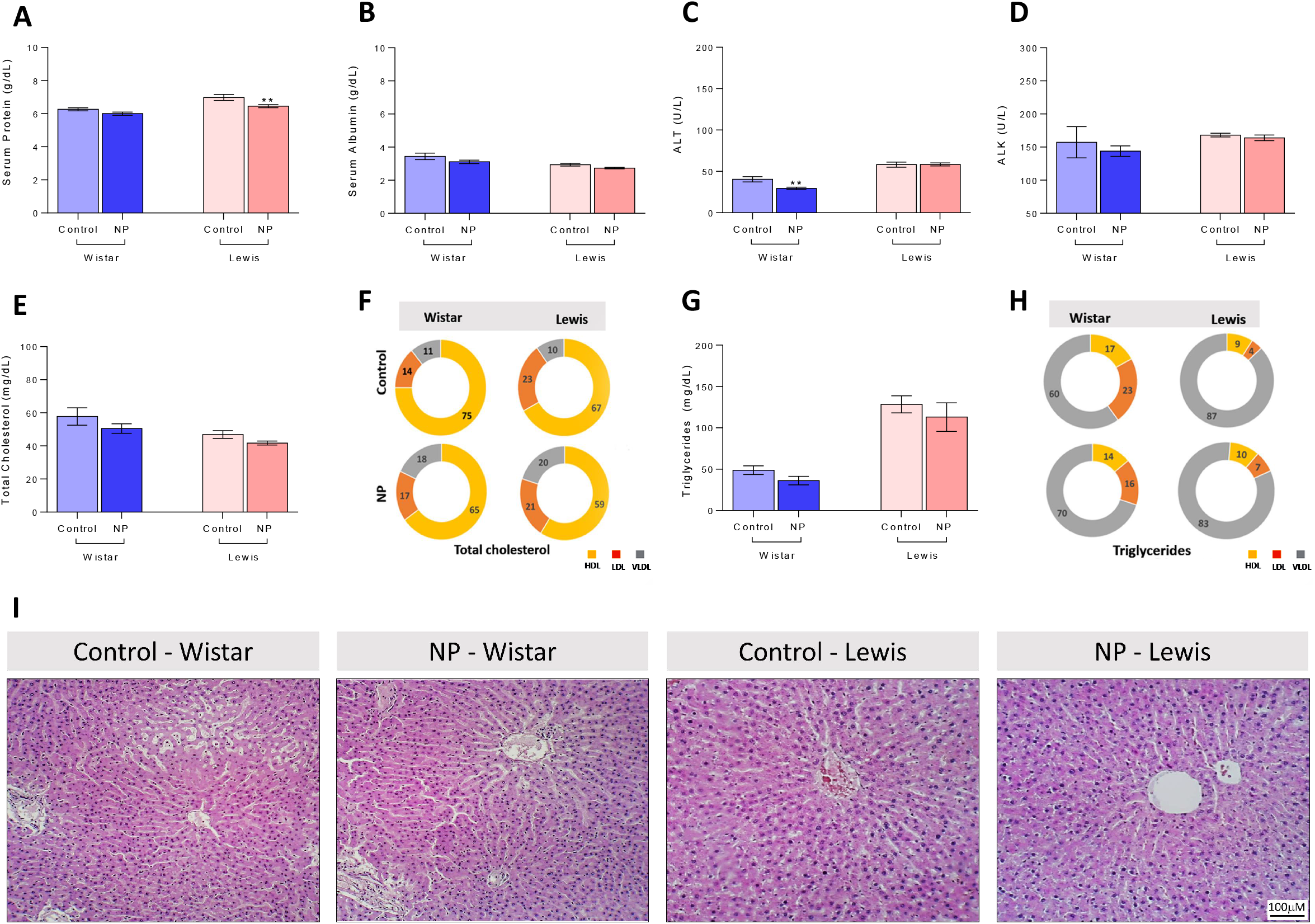
Hepatic Function: Serum **A)** Protein (g/dL), **B)** Albumin (g/dL), **C)** ALT (U/L), **D)** ALK (U/L), **E)** Total cholesterol (mg/dL), **F)** Cholesterol fractions (%), **G)** Total triglycerides (mg/dL) and **H)** Triglyceride fractions (%). **I)** Illustrative microphotographs of HE-stained liver sections under final 100x magnification.

**Figure 2.**
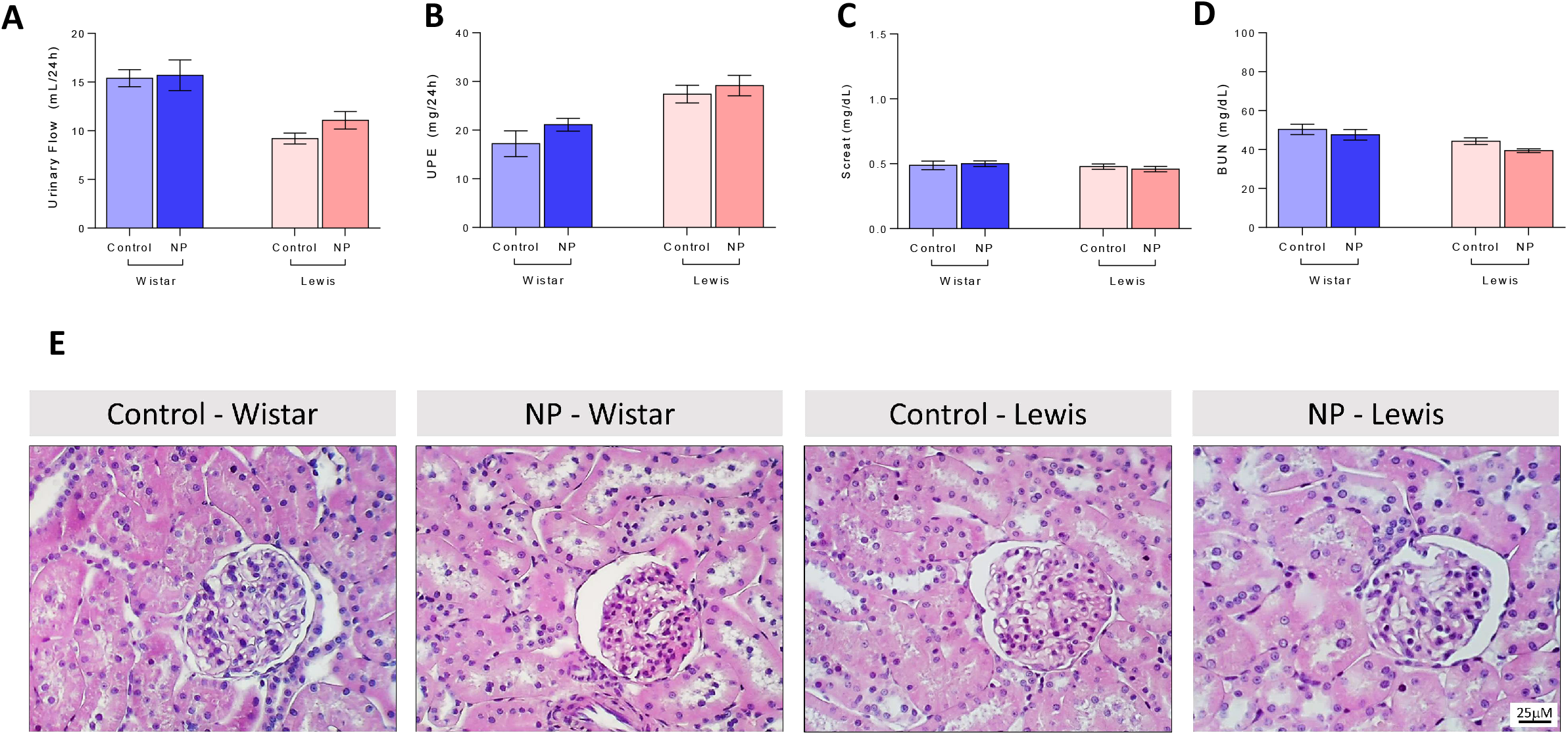
Renal Function: **A)** Urinary flow (mL/24h) **B)** Proteinuria (mg/24h), **C)** BUN (mg/dL), **D)** Screat (mg/dL). **E)** Illustrative microphotographs of HE-stained kidney sections under final 400x magnification.

Additional biomarkers of inflammation and overactivity of the coagulation system, usually correlated with worse prognosis in human patients infected by SARS-CoV-2, where also studied in NP animals. As shown in **Table 2**, both Wistar and Lewis rats receiving IM injections of NP exhibited regular values of serum CRP, Tnl and D-dimer, compared to Control animals.

**Table 2.**
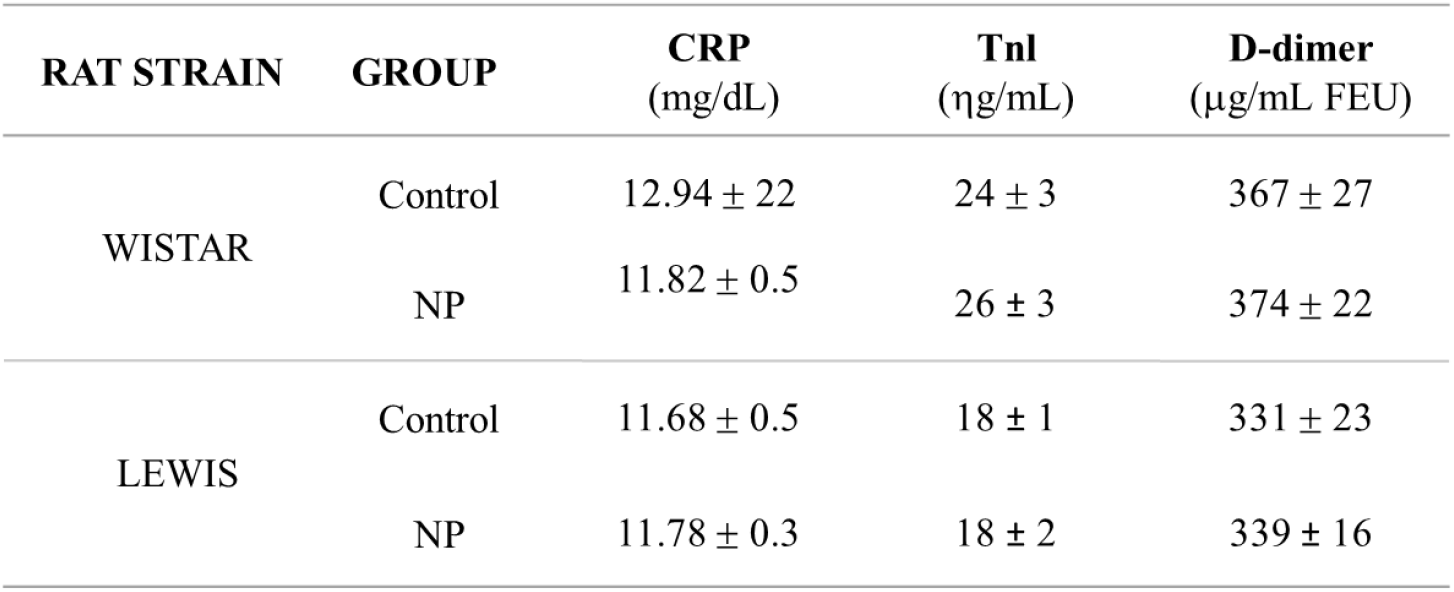
Indirect serum biomarkers of COVID-19: C-reactive protein (CRP, mg/dL), Troponin I (TnI, ng/mL) and D-dimer (μg/mL FEU) evaluated in the animals of each experimental group.

### NP injection induced specific immune response in both Wistar and Lewis rats, thus promoting the synthesis of specific antibodies

The presence of specific anti-SARS-CoV2 IgG in the serum of animals submitted to NP injections was evaluated by WB, as illustrated in **Figure 3A**. Each number at X axis of the barghraph represents one of the experimental animals. Positive dark bands at the point on the WB strip correspondent to the NP molecular weight (45KDa) indicate the positivity of serum sample for specific anti-NP IgG, which was evident in all NP Lewis animals, and in at least 5 of the 7 Wistar NP rats, indicating that the serum of these animals contained high levels of specific IgG anti-SARS-CoV2. Additionally, although the serum of the remaining 2 Wistar animals (rats 22 and 24) sensitized with NP showed only weak positivity when observed by naked eyes, it presented detectable positivity when analyzed with the Image Lab software (Bio-Rad), as can be seen in the correspondent bars above of each WB band photograph. Accordingly, none of the sera from Control Wistar or Lewis animals exhibited positive WB bands, indicating the absence of detectable anti-SARS-CoV2 antibodies in these groups, in spite of the similar concentrations of total serum globulins in NP animals compared to their strain-matched controls (**Figure 3B**). This observation becomes even more evident when WB band densities were statistically analyzed, as shown in **Figure 3C**. As observed in the respective bargraph, both Wistar and Lewis animals sensitized with the recombinant NP exhibited significantly high levels of anti-SARS-CoV2 IgG, compared to the Control groups, in which there were no crossreactivity.

**Figure 3.**
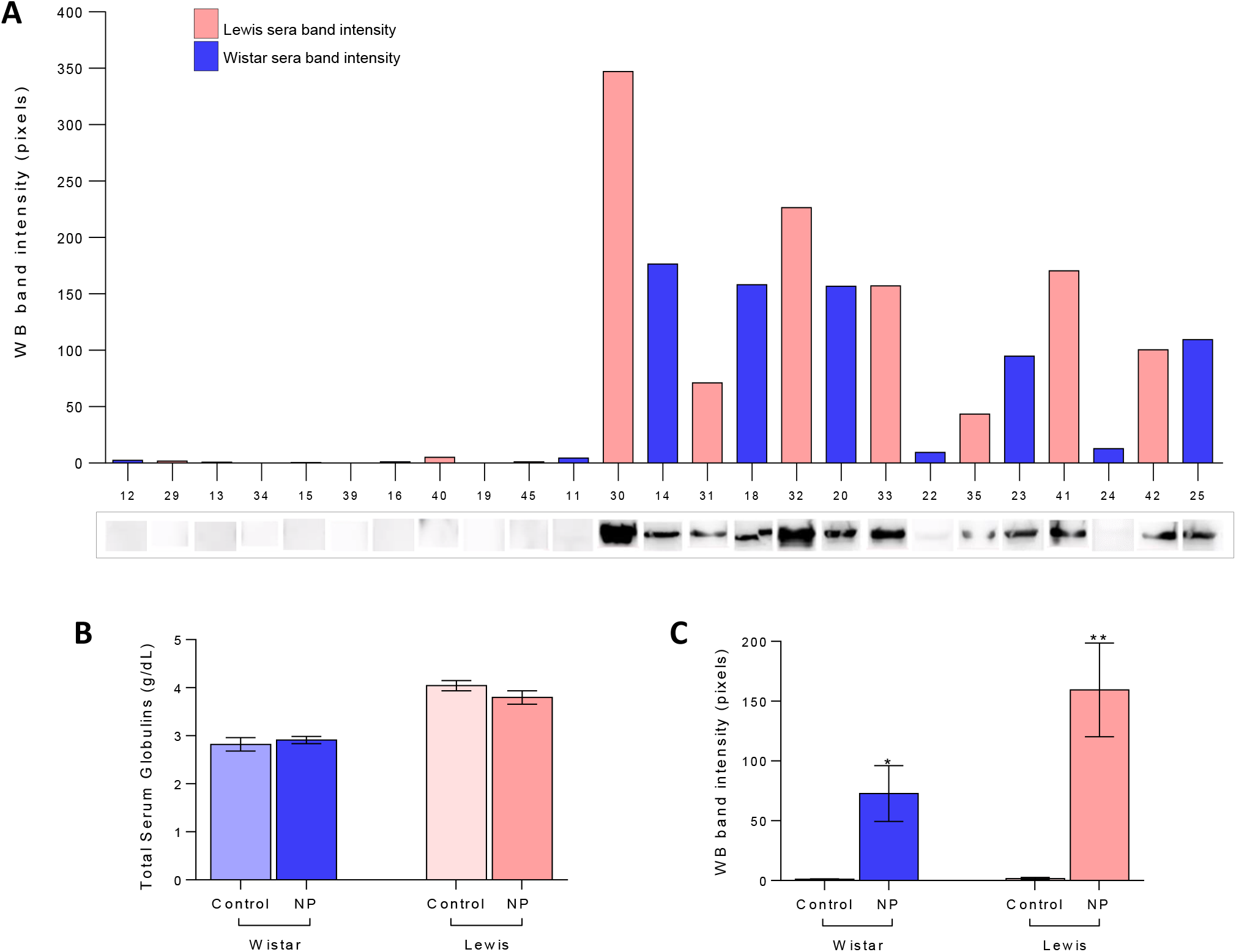
Immunization: **A)** Illustrative WB bands, pointing the presence of anti-NP specific IgG antibodies in the serum samples of both Wistar and Lewis rats submitted to NP inoculations **B)** Total serum globulin concentration (g/dL), **C)** WB band intensity quantification.

### NP IM injections induced the activation of the Bronchus-Associated Lymphoid Tissue (BALT) in both Wistar and Lewis rats

Histological analysis of lung tissue from NP and Control Wistar and Lewis rats were performed in HE-stained sections, by an experienced pathologist. As demonstrated in **Figure 4**, the lung parenchyma of Control rats (**A**, **C**) presented preserved appearance, regular alveolar septa and vessels. On the other hand, the lung of animals submitted to NP inoculation (**B**, **D**) exhibited altered pulmonary histoarchitecture, alveolar septa thickening and cellular infiltrate (**arrows**), which could be better seen under higher magnification in **J** and **L**. Moreover, NP animals also exhibited follicular bronchiolitis (**F**, **H**) accompanied by lymphocytic infiltrate (**arrows**), compared to its strain-matched controls. A bargraph representation of the score of inflammatory cells observed in the lung of NP and Control animals of each strain can be seen in **Figure 4U**. According to our results, both Wistar and Lewis NP animals exhibited significant increased lung cell infiltration, compared to Control groups. Further immunohistochemistry analysis was performed to better characterize the inflammatory cells infiltrating the lung tissue of the experimental animals. Pinkish-red points in the illustrative microphotographs of pulmonary tissue, shown in **Figure 4 M-P** are CD68+ macrophages. As can be noticed, Both Wistar and Lewis NP rats exhibited increased lung macrophage infiltration, compared to its strain-matched controls, however, only in Lewis NP animals this difference was statistically significant (**Figure 4V**). Conversely, brown points in the illustrative microphotographs of pulmonary tissue, shown in **Figure 4 Q-T** are CD3+ T-cells. Once more, Lewis NP rats exhibited increased lung lymphocyte infiltration, compared to its strain-matched however, this difference was not statistically significant (**Figure 4X**). An illustrative heat map was presented in **Figure 4Z**.

**Figure 4.**
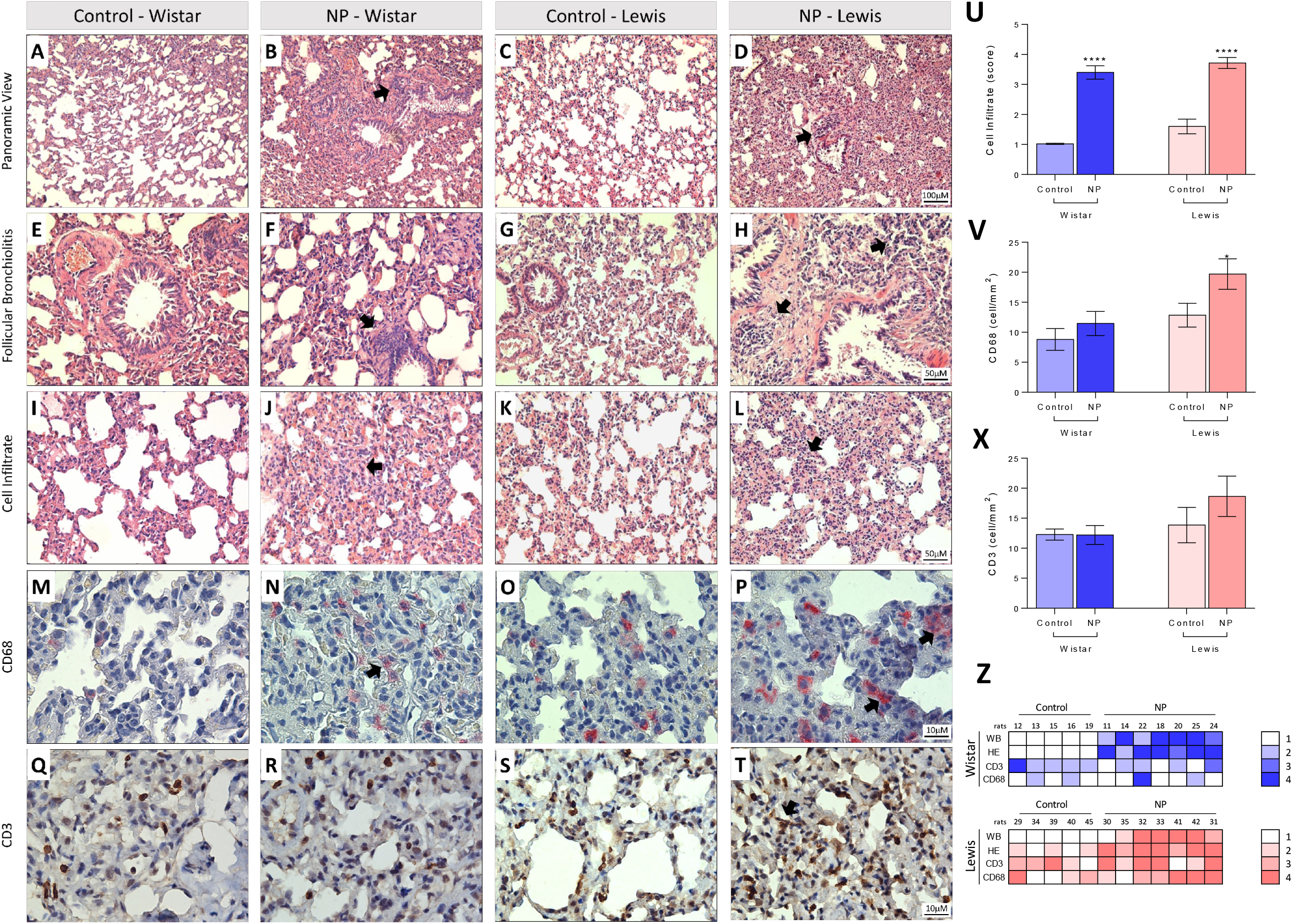
Histological assessment and immunohistochemistry in lung tissue stained with HE **(A-L)** and immunostaining by CD68 and CD3 **(M-T)**. In **A** and **C** the lung parenchyma presents normal appearance, as well as regular alveolar septa and vessels in the Wistar and Lewis Control group. In contrast, in **B** and **D** we identified distortion of pulmonary histoarchitecture and intense thickening in the alveolar septa region with the presence of cellular infiltrate and collagen deposition around the septa in the NP Wistar and Lewis groups (Arrows). Note in **F** and **H** the follicular bronchiolitis with the presence of intense lymphocytic infiltrate in the NP Wistar and Lewis groups when compared to the respective controls (**E** and **G**). Observe in **J** and **L** the thickening of pulmonary interstitium of the NP Wistar and Lewis groups (arrows) when compared to the controls group (**I** and **K**). In **N** we identified in pinkish red the immunostaining for CD68 in the pulmonary interstitium of the NP Wistar group, showing a slight number of cells positive for CD68 and when compared with its respective control **(M)**. In contrast, we observed an increase in cells immunostained for CD68 in lung parenchyma of NP Lewis group **(P)** when compared to its control group **(O)**. In **R**, note in brown the immunostaining for CD3 lymphocyte in the lung parenchyma NP Wistar group, when compared to the control group **(Q)** and in **T**, we identified CD68-positive cells in the thickening of pulmonary interstitium and control **(S)**. In **(U)**, graphical representation of the score for inflammatory cells indicate significant increase in NP Lewis and Wistar groups. In **(V)** observe CD68 statistical difference in Lewis NP and its control significant expression in alveolar septa and peribronchial region and **(X)** Graphical representation of CD3 quantification of CD3 in alveolar septa and peribronchial region of lung tissue. In **Z**, note intensity correlation of the anti-NP antibodies and expression of CD68 and CD3 in lung tissue of NP Wistar and Lewis group.

In order to verify whether the animals which exhibited a more prominent humoral immunological response against NP (with higher anti-NP IgG sera concentrations) would also be those who exhibited more exuberant lung histological alterations, a heat map was presented to correlate the production of anti-NP IgG, and immunohistochemistry in lung tissue stained with HE and immunostaining by CD68+ and CD3+ (**Figure 4Z**). As can be seen, a positive correlation between antibody production and immunohistochemistry in lung tissue stained with HE is observed mainly for Lewis rats with a correlation coefficient (r) of 0.8 than with Wistar (r of 0.3).

## DISCUSSION

SARS-CoV-2 is the causative agent of the 21^st^ century pandemic that has already killed more than half a million people worldwide. This virus still threatens public health worldwide with the emergence of new variants capable of being more transmissible, lethal, and reducing the efficiency of vaccines available on the market [20,21]. To date, experimental, pre-clinical and clinical studies are engaged to elucidate the pathophysiological mechanisms involved in COVID-19 progression and to seek for effective therapeutic strategies to avoid the development of severe manifestations and irreversible sequelae, especially in the most vulnerable patients. Additionally, since January 2020, the global scientific community is focused on developing and producing immunizing agents and vaccines to limit the spread and the gravity of COVID-19. Currently, the global efforts are concentrated on massive COVID-19 immunization, through both the distribution of the present vaccines for as many people all over the world, and through the stimulus to the local production of different immunizing agents in as many countries it is possible [11, 22]. In this context, the present study aimed to evaluate the impacts of the application of the recombinant SARS-CoV-2 NP, in 2 different rat strains, in order to verify if this inoculation would result in the development of specific immune response against SARS-CoV-2, represented by the presence of specific anti-NP IgG in the sera of injected animals. Additionally, we sought to verify the safety of NP as a potential immunizing agent, through the analyzes of the systemic effects of NP application on experimental animals.

No mortality, nor general behavioral alterations, inappetence or pain sings were observed in the animals of the different groups included in the present study. Vaccines usually produce only subtle clinical side effects, such as local pain due to the infiltration of inflammatory cells at the administration site, and slight fever, reflecting the expected immunological response of the host to the vaccine, similarly to what occurs after a natural infection These symptoms derive from the release of proinflammatory mediators such as the CRP, interleukins and coagulation factors, and are intrinsic of the activation of immune response [24]. Although vaccines are rarely related to systemic and local toxicity, due to the small amount of biological material which is injected for immunization, general health tests are mandatory when developing a new drug or treatment aimed for future clinical application. Accordingly, here we demonstrated that NP injections did not cause biochemical or histopathological changes which may suggest hepatotoxicity (increased concentration of liver enzymes, body weight loss, decreased serum protein concentration, dyslipidemia), or nephrotoxicity (polyuria/oliguria, proteinuria, increased Screat and BUN concentrations), and did not increased the main serum biomarkers of inflammation: CRP, TnI or D-dimer, thus providing satisfactory evidence that NP was safe for rats, exhibiting no toxic effects at the dose and route of administration tested in the experimental animals of this study.

As widely known, once a superior organism is exposed to invading microorganisms, such as viruses or bacteria, small parts of the pathogen stimulates the B-cells of the host, which in turn, differentiate into plasmocytes and start producing specific antibodies. In a very simplified way, antibodies are proteins able to recognizing and bind to known invading microorganisms, promoting both its neutralization or the opsonization of its phagocytosis by macrophages and neutrophils of the innate immune system of the host. Therefore, once an animal is sensitized to a particular pathogen, the response of its immune system in a future infection (reinfection with the same pathogen) will be faster and more effective, thus preventing the development of a more severe presentation of the disease [22, 23]. In order to evaluate the efficiency of NP to activate the immune response of inoculated animals, we analyzed the presence of specific anti-SARS-CoV2 IgG antibodies in the sera of NP-injected and non-injected rats, using WB. As expected, only sera from animals of the NP groups (both Wistar and Lewis strains) exhibited positivity for anti-SARS-CoV2 class G antibodies, therefore; it seems correct to state that the inoculation with recombinant NP in this experimental protocol succeeded in promoting the sensitization of the animals, with consequent synthesis of specific antibodies able of recognizing SARS-CoV2. Our results suggest the potential use of the recombinant NP described here, as both an immunizing agent and a useful tool for the production of rapid indirect diagnostic tests for SARS-CoV2, since WB strips containing this immobilized protein could be used to recognize sera from patients that have already been exposed to the virus, or have already had COVID-19 vaccine, and therefore would have specific strand-sensitive IgGs. In accordance with our findings, recent studies of Smits and collaborators comparing the specificity of IgG antibody response against SARS-CoV-2 demonstrated that, most of the COVID-19 patients exhibited specific immune response against NP, while few or none immunoreactivity against SP or MP have been found [25].

Finally, we analyzed lung samples of the animals of each experimental group, and we found that, recombinant NP IM injections triggered a local inflammatory reaction in the lung parenchyma of injected animals. Areas of multifocal expansion, as well as increased number of mononuclear cells, mainly lymphocytes, and macrophages, were noticed in the lungs of Wistar rats and, specially, in those from Lewis animals. Our histological and immunohistochemical analysis revealed that the greatest increase in lung infiltrating cells was predominantly within parenchymal regions rather than vascular areas and that it was mostly composed by macrophages and lymphocytes. Here, we report for the first time results describing pulmonary immune responses induced by the intramuscular injection of a recombinant SARS-CoV-2 protein. Our experimental findings provide data for further studies to better understand the effects of immunization against SARS-CoV-2, specifically on the lung parenchyma, since it is only possible with the use of laboratory animals. However, one of the limitations of using experimental animals are the inherent difficulties due to anatomic and physiological differences between other mammalian species and the human beings.

It is of note that the lower respiratory tract of mammals is constantly exposed to multiple airborne pathogens, and a prompt and effective immune response against such inhaled invaders is crucial for the survival of species [26, PABST 2010]. In this context, during the evolutionary process, a mucosal secondary lymphoid tissue, embedded in the walls of the large airways, the Bronchus-Associated Lymphoid Tissue (BALT) became part of the immune defense arsenal of many mammals. BALT is constitutive during all life phases in some species, including rats and rabbits, but it is absent in healthy adult mice and humans [Sminia et al., 1989, Troy D. Randall 2010]. Although more studies with different species, more similar to the human beings, regarding the specific pulmonary immune response are still required, we can suppose that, interestingly, the IM immunization with recombinant NP was able to activate the BALT of the rats, as a response of the systemic immune response triggered by the immunization, reflecting in the overactivity of immune cells observed in the lung of NP rats. Our findings suggest that, in spite of the IM administration, our immunizing agent was able to trigger an immune response specifically in the lungs of the injected rats. This is clinically relevant: the immunomodulation being a primary mechanism underlying COVID-19 supports the prioritization of tolerance as a vaccine strategy.

## CONCLUSIONS

The data presented in this manuscript have several implications for our understanding on the cellular and humoral mechanisms triggered by SARS-CoV-2 immunizing components. Based on our results, we can conclude that the inoculation of recombinant NP protein in Wistar and Lewis rats was safe, since it did not promote changes in the main clinical, biochemical and histological parameters studied here, which could indicate toxicity of this compound. Additionally, NP inoculation successfully promoted the production of specific IgG antibodies against SARS-CoV2, indicating its potential use for massive vaccination immunization, especially because it seems to induce specific pulmonary protection, although additional studies are still needed.

## Funding Support

Financial and material support was provided by the São Paulo Research Foundation (FAPESP) grants, as follows: (FAPESP #2018/20403-6 for author VLC, FAPESP #2019/00195-2 and FAPESP #2020/04680-0 for author CRG, FAPESP #2019/21739-0 for author CGB, FAPESP #2018/00415-5 for author WPRT and #2019/19939-1 for author ICS) and by Brazilian Government Research Funding Foundation (FINEP) MCTI Rede Virus grant 0459/20 for author CRG. The author APB was supported by the Coordination of Superior Level Staff Improvement (CAPES) grant 88887.508606/2020-00.

## Acknowledgments

We would like to thank FAPESP and FINEP for financial support and Wiener laboratories by performing CRP, D-dimer and TnI serum dosages.

## Notes

### Competing Interest Statement

The authors have declared no competing interest.

